# Carbapenem-Resistant *Enterobacter hormaechei* Uses Mucus Metabolism to Facilitate Gastrointestinal Colonization

**DOI:** 10.1101/2024.09.25.615021

**Authors:** Ritam Sinha, Elizabeth N. Ottosen, Tshegofatso Ngwaga, Stephanie R. Shames, Victor J. DiRita

## Abstract

The emergence and global spread of carbapenem-resistant *Enterobacter cloacae* complex species presents a pressing public health challenge. Carbapenem-resistant *Enterobacter* species cause a wide variety of infections, including septic shock fatalities in newborns and immunocompromised adults. The intestine may be a major reservoir for these resistant strains, either by facilitating contamination of fomites and transfer to susceptible individuals, or through translocation from the gut to the bloodstream. For this reason, we sought to establish a neonatal mouse model to investigate the mechanisms underpinning gut colonization by carbapenem-resistant *Enterobacter hormaechei*. We describe a new mouse model to study gut colonization by *Enterobacter* species, leading to vital insights into the adaptation of carbapenem-resistant *E. hormaechei* to the gut environment during the early stages of intestinal colonization. We observed successful colonization and proliferation of *E. hormaechei* in the five-day old infant mouse gut, with primary localization to the colon following oral inoculation. We also uncovered evidence that *E. hormaechei* uses mucus as a carbon source during colonization of the colon. Our findings underscore the importance of oxygen-dependent metabolic pathways, including the pyruvate dehydrogenase complex, and *N-*acetyl-D-glucosamine metabolism, in gut colonization and proliferation, which aligns with previous human studies. These insights are essential for developing novel therapeutic strategies that can serve as decolonization therapies in at-risk populations.

**Importance:** Bloodstream infections caused by *Enterobacter* species pose a significant clinical threat. The intestine acts as the primary site for colonization and serves as a reservoir for infection. To combat this pathogen, it is crucial to understand how carbapenem-resistant *Enterobacter* species colonize the gut, as such knowledge can pave the way for alternative therapeutic targets. In this study, we developed a novel neonatal mouse model for gastrointestinal colonization by *Enterobacter* species and discovered that mucus plays a key role as a carbon source during colonization. Additionally, we identified two mucus catabolism pathways that contribute to intestinal colonization by carbapenem-resistant *E. hormaechei*. This new mouse model offers valuable insights into host-pathogen interactions and helps identify critical gastrointestinal fitness factors of *Enterobacter*, potentially guiding the development of vaccines and alternative therapeutic strategies to minimize intestinal carriage in patient populations at risk for infection with *Enterobacter* species.

## Introduction

The emergence and spread of extensively antibiotic-resistant Gram-negative bacteria, specifically carbapenem-resistant *Enterobacteriaceae* (CRE) poses a significant and urgent global health challenge (1, 2). Infections caused by CRE are major contributors to illness and death, with children and the elderly particularly affected on a global scale (3). CRE infections commonly lead to sepsis, notably in hospitalized immunocompromised patients, neonates, and preterm infants. Emergent pathogens from the *Enterobacter cloacae* complex (ECC) contribute to septic shock fatalities in newborns, termed late-onset neonatal sepsis, as well as in immunocompromised adults (4). The ECC, comprised of multiple species, poses identification challenges at the species level, resulting in possible misidentification as *Enterobacter cloacae*. Recently, *Enterobacter bugandensis* and *Enterobacter hormaechei*, two species within the ECC, have been identified in cases of neonatal sepsis (5, 6). However, the precise molecular mechanisms of *Enterobacter* pathogenicity, including infection routes and virulence factors, remain inadequately defined.

Bloodstream infections caused by CRE predominantly originate from hospital-associated sources like contaminated fomites (5). Additionally, the intestine serves as a primary site of colonization and reservoir for infection. Intestinal colonization not only facilitates spread to fomites in the clinical setting, but an emerging hypothesis posits that colonizing CRE strains can translocate from the gut to the bloodstream under certain conditions, such as during antibiotic treatment. One study demonstrated a correlation between mortality from neonatal sepsis, gut colonization by CRE strains, and broad-spectrum antibiotic treatment (7), and that neonatal gut colonization or carriage of CRE strains is linked to the mother’s microbiota. Consequently, there is an urgent need for alternative therapeutic strategies to prevent CRE colonization of the gut, or to decolonize at-risk patients who are already colonized with CRE. To achieve this goal, it is imperative to understand the mechanisms facilitating gut colonization by *Enterobacter* species, and this in turn requires a robust animal model.

Adult mice with a healthy gut microbiota naturally prevent colonization with various pathogenic enteric bacteria. Consequently, antibiotic treatment is essential to facilitate colonization in the laboratory. Several adult mouse models have been developed to study colonization by CRE strains (8, 9). These require treatment with sodium bicarbonate to neutralize stomach acid, and/or antibiotic treatment to reduce the normal gut microbiota. The objective of our current study was to develop a neonatal mouse model to elucidate mechanisms by which *Enterobacter* species establish gut colonization. Neonatal mice have a limited microbiota, thereby obviating treatments like sodium bicarbonate or antibiotics to establish gut colonization, and instead mirroring natural colonization in humans. Neonatal mice are widely used to study different enteric and opportunistic pathogens such as *Vibrio cholerae*, *Shigella,* ETEC, and *Klebsiella pneumoniae* (10–13). These mice also offer advantages for research given their ease of handling, cost-effectiveness, and amenability to genetic manipulation and drug treatments. For instance, manipulating factors like epidermal growth factor (EGF) allows for investigation of how maternal factors received from breast milk influence the translocation of gut-resident pathogenic bacteria across the intestinal barrier and to secondary systemic sites (14). Finally, as human neonates are at risk for neonatal sepsis caused by CRE, a neonatal mouse model is likely to more accurately replicate features of human infection.

We sought to understand mechanisms by which *Enterobacter* species adapt to the gut environment and establish colonization. We demonstrate that these strains efficiently colonize the infant gut, primarily the colon, during the initial phase of intestinal colonization. Our work also suggests that mucus is a vital carbon source facilitating bacterial survival and proliferation in this environment. Our findings also highlight the significance of oxygen-dependent metabolic pathways (such as the pyruvate dehydrogenase complex and *N-*acetyl-D-glucosamine metabolism), and mucin metabolism during *E. hormaechei* gut colonization. This study provides insights into how carbapenem-resistant *E. hormaechei* adapts to the gut environment during the initial phase of colonization, offering valuable knowledge to develop alternative therapeutic approaches.

## Results and Discussion

### *Enterobacter* species efficiently colonize the GI tract of infant mice

We aimed to develop a neonatal mouse model to identify mechanisms of early-stage gut colonization by *Enterobacter* species. For this, we characterized the ability of different *Enterobacter* strains to colonize the intestinal tract of suckling mice. We selected three strains: *Enterobacter cloacae* ATCC_13047, and two carbapenem-resistant *Enterobacter* clinical isolates (CRE13 and CRE14). To assess gut colonization by these strains, we inoculated five-day-old CD-1 mice with 10^5^ colony forming units (CFU) and measured the bacterial load in the entire intestine after 24 hours. To avoid potential inhibition of *Enterobacteriaceae* colonization by maternal breast milk (14), we separated infants from the dams after inoculation. Bacterial burden of these strains in the intestinal tract ranged from 10^5^ to 10^7^ CFU per gram of tissue (**Figure 1A**). We then characterized population dynamics of the three *Enterobacter* strains along the length of the intestinal tract 24 hours post-inoculation. We observed the highest bacterial burden in the colonic portion of the intestinal tract for all three *Enterobacter* strains, ranging from 10^7^ to 10^8^ CFU per gram of tissue (**Figure 1B**). Only CRE13 was detected at a low level (10^4^ CFU/g tissue) in the distal portion of the small intestine 24 hours post-inoculation, whereas ATCC_13047 and CRE14 were below the limit of detection in this site. These data indicate that *Enterobacter* species colonize the intestinal tract of infant mice and primarily localize to the colon, similar to observations from a neonatal mouse model of *Klebsiella pneumoniae* colonization (13). For the remainder of this study, we focused on *E. hormaechei* CRE14 strain because in separate works we have investigated its growth dynamics and fitness factors during bloodstream infection in an adult mouse model (15, 16).

**Figure 1.**
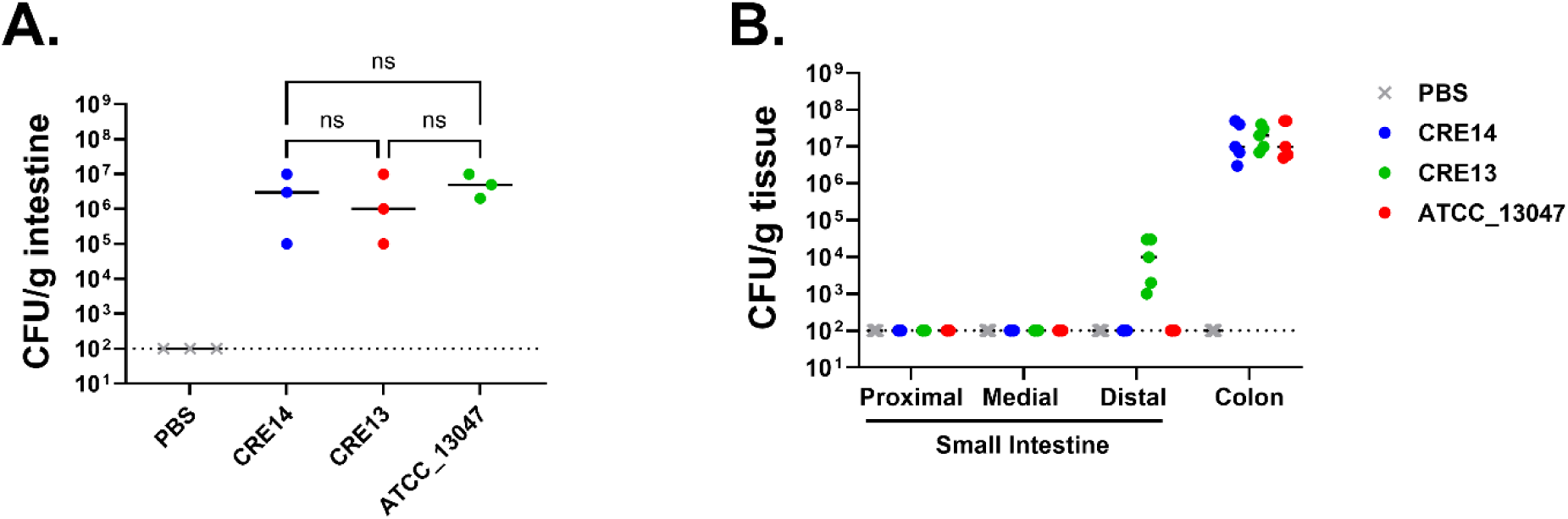
Colonization of the infant mouse gut by different *Enterobacter* species isolates: A) Recovery of three *Enterobacter* strains, including two carbapenem-resistant clinical isolates (CRE14 and CRE13) and *Enterobacter cloacae* ATCC_13047, from the entire intestinal tract of 5-day-old CD-1 mice 24 hours post-inoculation with 10^5^ CFU orally (n=3 mice per group). Statistical significance was determined by One-way ANOVA with multiple comparisons. B) Recovery of *Enterobacter* strains from the small intestine (proximal, medial, distal) and colons of 5-day-old CD-1 mice 24 hours post-inoculation with 10^5^ CFU orally (n=5 mice per group). Each point represents the bacterial burden recovered from a single mouse and horizontal lines are the median CFU/g of tissue of each group. Dashed lines represent the limit of detection.

Varying disease susceptibility between inbred and outbred infant mice following enterotoxigenic *E. coli* infection has been observed (17), so we investigated whether asymptomatic colonization by *E. hormaechei* could also vary based on mouse strain. To assess this, we selected one outbred (CD-1) and two inbred (C57BL/6 and BALB/c) mouse strains. Mice were orally inoculated with 10^5^ CFU of CRE14, and bacterial burden in the colon was determined 24 hours post-inoculation. We observed that CRE14 colonized the colons of different mouse strains to approximately the same level, 10^7^ to 10^8^ CFU per gram of tissue, suggesting that this model can be applied to a variety of mouse strains and highlighting its versatility (**Figure 2A**). To assess how dose impacts CRE14 intestinal colonization, we selected four different doses (10^3^, 10^5^, 10^7^, and 10^8^ CFU) and orally administered these to CD-1 pups. Twenty-four hours post-inoculation, we observed poor colonization in the colons of mice inoculated with 10^3^ CFU dose of CRE14. The bacterial load was below the limit of detection in three out of five mice, while we recovered approximately 10^5^ CFU per gram of tissue from two animals. Significantly higher burdens ranging from 10^7^ to 10^9^ CFU per gram of tissue were observed in the colon following infection with higher doses (10^5^ to 10^8^ CFU) (**Figure 2B**). The presence of an endogenous *Enterobacteriaceae* species in adult mice hinders *Salmonella* Typhimurium colonization, particularly when administered at lower doses (18). In the CD-1 infant mouse gut, we identified an endogenous lactose fermenting *Enterobacteriaceae* isolate sensitive to ampicillin. As CRE14 is a very weak lactose fermenter, this endogenous strain was easily distinguished from CRE14 by plating on MacConkey agar. The endogenous strain was present to the same level in the colons of PBS and CRE14 inoculated mice (**Supplemental Figure 1A**). This finding suggests that at higher doses, CRE14 can effectively compete with this endogenous *Enterobacteriaceae* to co-colonize the colon. Finally, at the highest dose of 10^8^ CFU per mouse, we recovered CRE14 from the colon and also from deeper tissue such as the liver and spleen in three out of five mice (**Figure 2B**). Systemic infection did not cause mortality in pups by our 24-hour timepoint. These data indicate that at higher oral doses, CRE14 breaches the gut barrier, disseminating to secondary sites.

**Figure 2.**
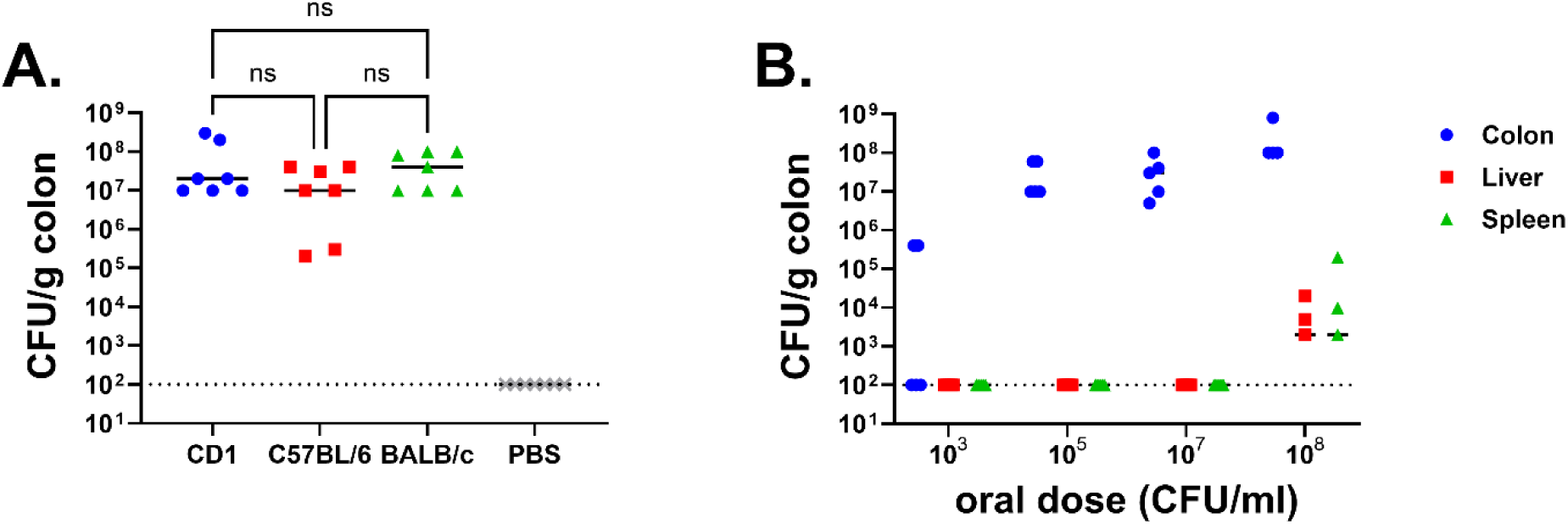
Colonization ability of carbapenem-resistant *Enterobacter hormaechei* in different strains of mice: A) Recovery of CRE14 from the colons of three different strains of infant mice: CD-1 (outbred), C57BL/6, and BALB/c (inbred) 24 hours post-inoculation with 10^5^ CFU orally (n=7 mice per group). Statistical significance was determined by One-way ANOVA with multiple comparisons. B) CRE14 colonization in different organs (colon, liver, and spleen) in CD-1 infant mice 24 hours post-inoculation with varying oral doses (10^3^, 10^5^, 10^7^, and 10^8^ CFU) (n=5 mice per group). Each point represents the bacterial burden recovered from a single mouse, and horizontal lines are the median CFU/g colon from each group. Dashed lines represent the limit of detection.

Systemic spread from the gut to blood-filtering organs suggests damage to the gut epithelium. To visualize potential damage, we performed H&E staining on colon samples from mice inoculated with PBS or 10^8^ CFU of CRE14 (**Supplemental Figure 2A and B**). At high oral doses of CRE14 we detected mild inflammatory responses such as epithelial cell destruction and neutrophil infiltration (**Supplemental Figure 2C**). We also assessed the invasive potential of CRE14 using a gentamicin protection assay on human colon carcinoma cells (CaCo2) (19). At an MOI of 100 we found that CRE14 effectively adheres to CaCo2 cells, reaching approximately 10^7^ CFU/ml after a 30-minute incubation. Subsequently, we recovered approximately 10^5^ CFU/ml of CRE14 following treatment with gentamicin for two hours (**Supplemental Figure 2D**). These data suggest that invasiveness of CRE14 enables it to traverse from the gut to systemic organs, potentially leading to systemic infection, and suggests that this mouse model can be used to further investigate this process.

While we are primarily focused on initial colonization mechanisms of *Enterobacter* species, we also explored the feasibility of this model for studying longer term colonization. We orally administered 10^5^ CFU of CRE14 to pups, returned pups to their dams immediately after, and monitored bacterial burden in the colon for three days. Twenty-four hours post-inoculation we recovered up to 10^8^ CFU per gram of colon tissue. By day three, we detected 10^5^ to 10^8^ CFU per gram of colon tissue in five out of seven mice, while CRE14 fell below the limit of detection in two out of seven mice (**Figure 3A**). These findings suggest that *E. hormaechei* can rapidly colonize and proliferate in the infant gut shortly after inoculation, and this colonization persists up to three days post-inoculation.

**Figure 3.**
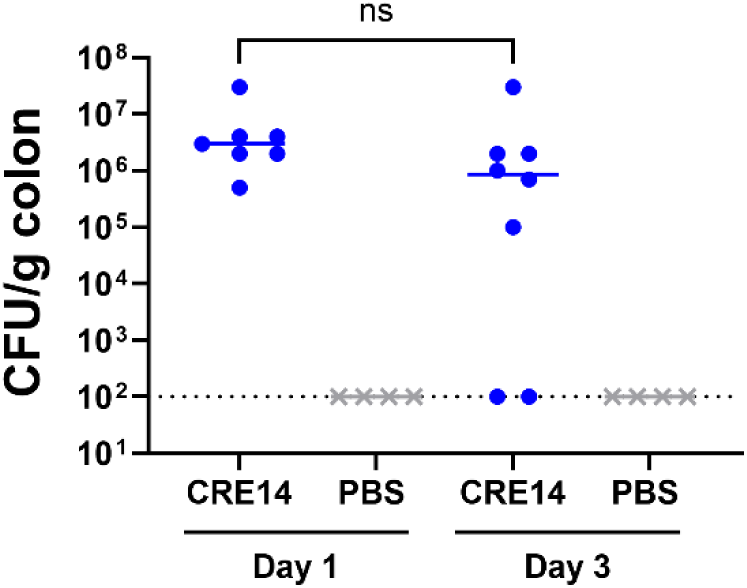
Durability of *E. hormaechei* gut colonization: Bacterial burden in the colons of 5-day-old CD-1 mice 1-and 3-days post-inoculation with 10^5^ CFU oral dose of CRE14 (n=7 mice per group). Each point represents the bacterial burden recovered from a single mouse. Horizontal lines represent the median CFU/g of colon tissue of each group. Statistical significance was determined by t-test. The dashed line represents the limit of detection.

A previous study emphasized the similarity in intestinal physiology development between full-term mice and premature human infants (20). Infant mice are born less developmentally mature than to-term human infants, and thus more closely resemble premature human infants. Additionally, an abundance of carbapenem-resistant *Enterobacter* species were observed in the intestinal tract of preterm infants in both developed and middle-income countries (3). Given these parallels, we propose that this neonatal mouse model could serve as a tool to explore mechanisms of how carbapenem-resistant *Enterobacter* species rapidly adapt to and establish asymptomatic carriage in the intestinal tracts of preterm human infants.

### Mucin serves as a potential carbon source for carbapenem-resistant *E. hormaechei* in the neonatal gut

To characterize the progression of CRE14 colonization, we measured bacterial burden in the colons of five-day-old CD-1 mice at different time intervals following oral inoculation with 10^5^ CFU. We recovered a small amount of CRE14 in the small intestine 2 hours post-inoculation, which fell below our limit of detection after 12 hours. Conversely, we observed an increase in bacterial burden in the colon over time, reaching approximately 10^8^ CFU per gram of tissue after 24 hours (**Figure 4A**). These data demonstrate that CRE14 primarily grows in the colon of infant mice over the course of 24 hours, and that it is well adapted to the infant gut environment and has access to nutrients supporting its growth. Pathogenic and commensal *E. coli* strains can use mucus as a carbon source in the mouse intestine (21, 22). *E. hormaechei* has been shown to use galactose and *N-*acetyl-D-glucosamine as carbon sources, which are key components of mucus (23). We hypothesized that mucus serves as a major carbon source during colonization of the neonatal mouse gut. To test the growth of *E. hormaechei* when mucus is the sole carbon source, we cultured CRE14 in M9 minimal media containing 0.5% partially purified porcine gastric mucin and measured growth by CFU count over time. We observed significant CRE14 growth in this medium, reaching 1000-fold increase in CFU by 24 hours (**Figure 4B**). These data suggest that *E. hormaechei* can use mucin-derived sugars as a carbon source. This is consistent with a previous study that found certain *Proteobacteria* including *Klebsiella*, *Mixta*, *Serratia*, and *Enterobacter* encode several mucin-degrading glycosyl hydrolases (24).

**Figure 4.**
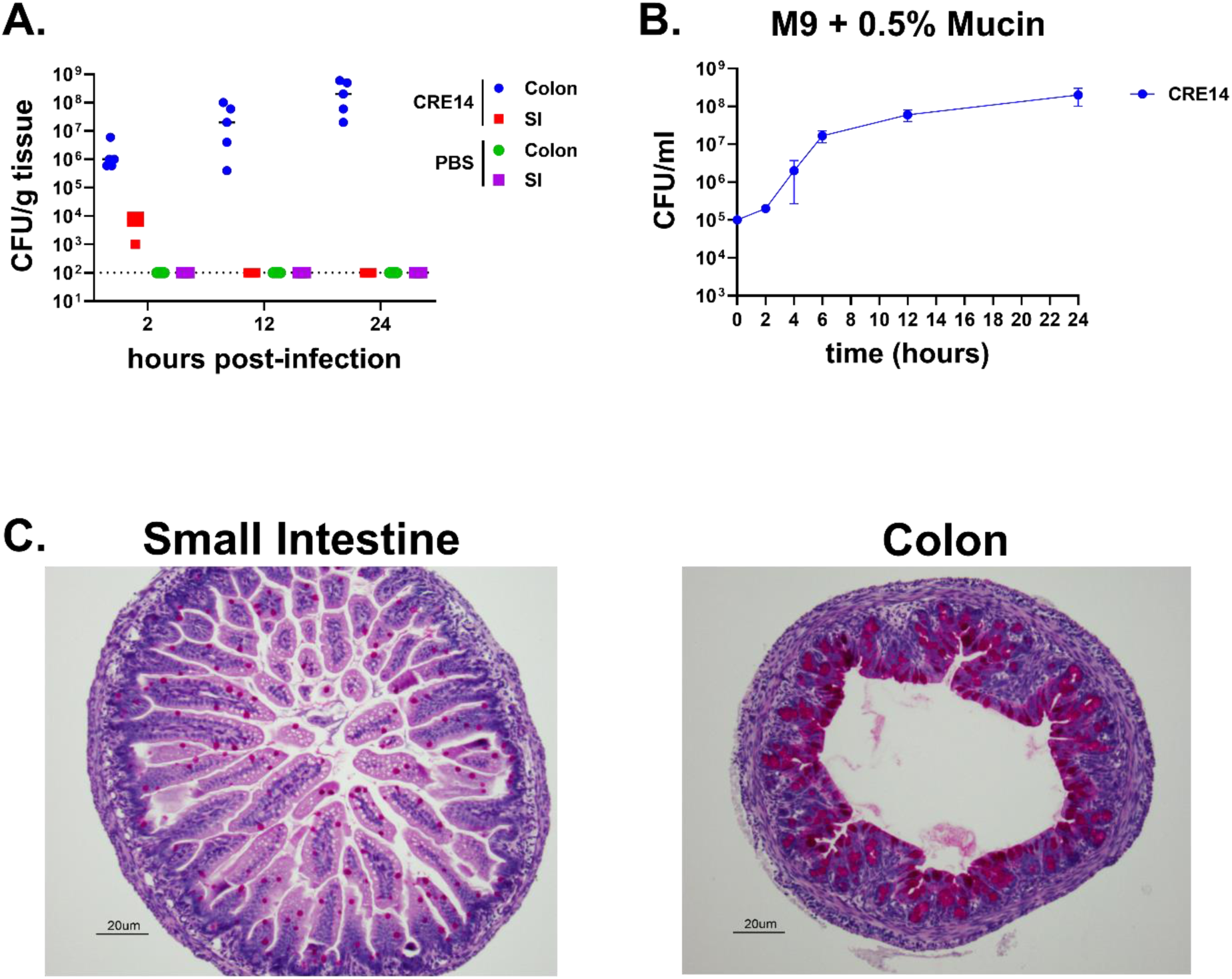
Rapid proliferation of carbapenem-resistant *E. hormaechei* in the colon of infant mice during short term colonization: A) Recovery of CRE14 from the colon and small intestine of 5-day-old infant mice 2-, 12-, and 24-hours post-inoculation with 10^5^ CFU orally (n=5 mice per group). Each point represents bacterial burden recovered from a single mouse. Horizontal lines represent the median CFU/g of tissue of each group. The dashed line indicates the limit of detection. B) Growth of CRE14 in M9 minimal media containing 0.5% partially purified porcine gastric mucin over 24 hours under aerobic conditions. Points represent the mean of three biological replicates and error bars are ± SEM. C) Abundance of mucin producing goblet cells in the small intestine and colon of 5-day-old infant mice determined by Periodic Acid-Schiff (PASH) staining. Representative images were selected from two independent experiments.

Mucus is unevenly distributed along the intestinal tract, with thicker mucus layers lining the stomach and colon, and thinner layers in the small intestine (25). We hypothesized that mucus availability could contribute to CRE14 localization to the infant mouse colon. To assess the abundance of mucus in the neonatal mouse gut, we performed mucin-specific Periodic Acid-Schiff (PASH) staining for carbohydrate compounds on samples from the distal portion of the small intestine and colons of un-inoculated infant mice. Staining was more abundant in the colon compared to the small intestine (**Figure 4C**), and we speculate that the higher abundance of mucin in the infant mouse colon could drive preferential *Enterobacter* colonization of the colon over the small intestine. Overall, these data indicate that mucin may serve as a highly abundant carbon source for *Enterobacter* growth during colonization of the infant gut.

### Carbapenem-resistant *E. hormaechei* growth on mucin is supported by the pyruvate dehydrogenase complex

To further explore growth of *E. hormaechei* in the intestine, we examined the pyruvate dehydrogenase complex (PDH). The PDH complex plays a significant role in growth of the pathogen *Vibrio cholerae* in the infant mouse gut (26), the genes are abundantly expressed by *E. coli* growing in the mouse colon (27) and it is a colonization factor for *Klebsiella* in the gut of adult mice (28). To examine the role of the pyruvate dehydrogenase complex in *E. hormaechei* growth on mucin and colonization of the infant mouse gut, we generated a strain lacking *aceE*, which encodes one subunit of the PDH complex. Initially, we characterized growth kinetics of the Δ*aceE* mutant in minimal media containing glucose as the only carbon source. Since the PDH complex converts pyruvate derived from glycolysis into acetyl-CoA to start the TCA cycle, we hypothesized the Δ*aceE* mutant would be unable to grow on glucose as the sole carbon source. We cultured wild-type and the Δ*aceE* mutant aerobically in M9 minimal media containing glucose and assessed growth by measuring optical density over time. Under aerobic conditions, where the PDH complex is active, the Δ*aceE* mutant is unable to grow when glucose is the only carbon source (**Supplemental Figure 3A**).

As *aceE* is the second gene in an operon (*pdhR-aceE-aceF*), we complemented Δ*aceE* using plasmid-encoded wild type *aceE* under control of the *pdhR* promoter. We then grew five strains – CRE14, CRE14 containing an empty plasmid (CRE14/vector), the Δ*aceE* mutant, Δ*aceE* with an empty plasmid (Δ*aceE*/vector), and Δ*aceE* complemented with the wild-type *aceE* allele (Δ*aceE/*p*aceE*) – in M9 minimal media containing glucose. While Δ*aceE* and Δ*aceE/*vector displayed a growth defect relative to wild-type CRE14, growth was partially restored to wild-type levels when the mutant was complemented with the wild-type *aceE* allele *in trans* (**Supplemental Figure 3B**). We hypothesized that full restoration to wild-type growth did not occur in the in the mutant complemented *in trans* because plasmid expression altered the wild-type stoichiometry of the large PDH complex, which has been seen previously (26, 29). To ensure the Δ*aceE* growth defect was due solely to the *aceE* mutation, we tested if we could restore its growth by supplementing the media, hypothesizing that adding both glucose and acetate to minimal media would restore Δ*aceE* growth to wild-type levels. *E. hormaechei* CRE14 encodes a phosphoenolpyruvate carboxylase (*ppc*), which converts phosphoenolpyruvate derived from glycolysis into the TCA intermediate oxaloacetate. Oxaloacetate and acetyl-CoA combine to make citrate which feeds into the TCA cycle. CRE14 also encodes an acetate-CoA ligase (*acs*), which can convert acetate to acetyl-CoA, thus mitigating loss of *aceE* and enabling the TCA cycle to proceed. To test this, we cultured CRE14 and the Δ*aceE* mutant in minimal media containing acetate alone, or glucose and acetate. Neither strain grew with acetate as the sole carbon source (**Supplemental Figure 3C**), and consistent with our reasoning, growth of the Δ*aceE* mutant was completely restored to wild-type levels when provided with both glucose (to produce oxaloacetate) and acetate (to produce acetyl Co-A) (**Supplemental Figure 3D**). These data indicate that the observed growth defect by the Δ*aceE* mutant is most likely due only to loss of *aceE* functionality. Click or tap here to enter text.

To investigate the role of the PDH complex in CRE14 growth on mucin, we cultured wild-type and the Δ*aceE* mutant in M9 minimal media containing 0.5% partially purified porcine gastric mucin and measured growth by CFU count for 24 hours. The Δ*aceE* mutant exhibited slow growth compared to wild-type, but ultimately reached similar levels after 24 hours in mucin (**Figure 5A**). These data suggest that the PDH complex contributes to wild type growth of *E. hormaechei* on mucin as the sole carbon source; our data also suggest that the amino acids present in mucin also do not support wild type growth of the Δ*aceE* mutant. We tested directly whether amino acids could support growth of the mutant in otherwise minimal media and confirmed that they do not (**Figure 5F**).

**Figure 5.**
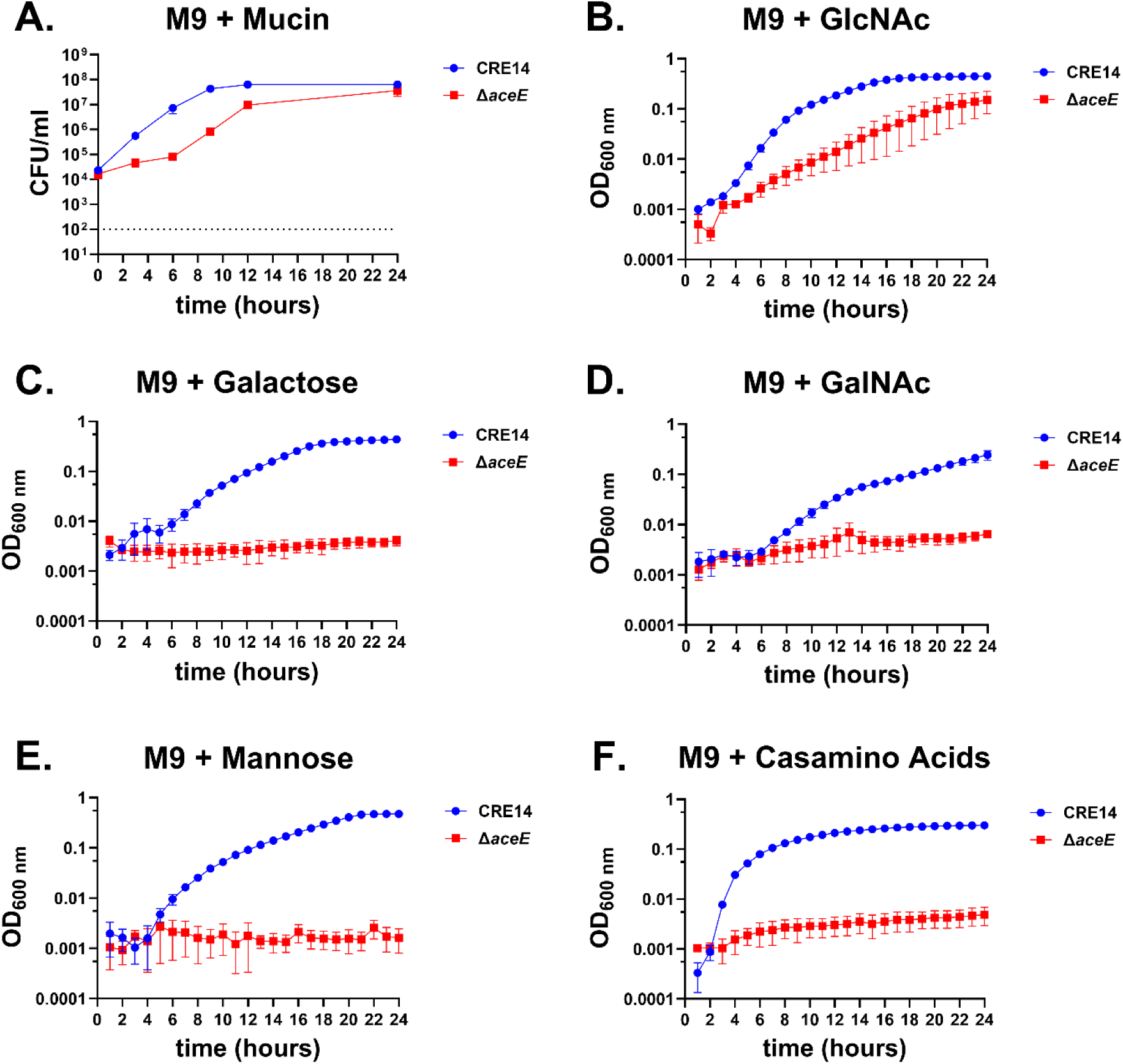
The *E. hormaechei* pyruvate dehydrogenase complex is essential for full growth in mucin: A) Growth of CRE14 and the Δ*aceE* mutant in M9 minimal media supplemented with 0.5% partially purified porcine gastric mucin over 24 hours. Data represent the mean CFU of three biological replicates and error bars represent the mean ± SEM. The dashed line indicates the limit of detection. B-F) Growth of WT and the Δ*aceE* mutant in M9 minimal media supplemented with 10mM glucose, *N-*acetyl-D-glucosamine (GlcNAc), galactose, *N-*acetyl-D-galactosamine (GalNAc), mannose, or 0.2% casamino acids over 24 hours. Data represent the mean OD_600nm_ of three biological replicates and error bars represent ± SEM.

MUC2, abundant in the colon, is composed of O-glycosylated and N-glycosylated carbohydrates. O-glycans primarily comprise branched carbohydrates such as *N*-acetyl-D-glucosamine (GlcNAc), *N-*acetyl-D-galactosamine (GalNAc), fucose, and sialic acid (30). In contrast, *N*-glycosylated carbohydrates are typically linked with mannose residues. To explore how CRE14 uses mucin-associated carbohydrates and to understand recovery of the Δ*aceE* mutant at later timepoints, we cultured wild-type CRE14 and the Δ*aceE* mutant in M9 minimal media containing GlcNAc, galactose, GalNAc, fucose, mannose, or sialic acid at 10 mM concentrations. CRE14 could not grow with fucose or sialic acid as carbon sources (**Supplemental Figure 4A & B**), but was able to grow with GlcNAc, galactose, GalNAc, or mannose as the sole carbon source (**Figure 5B, C, D, and E**). As anticipated, the Δ*aceE* mutant was unable to grow on galactose, GalNAc, and mannose. However, Δ*aceE* was able to grow, albeit slowly, in minimal media containing GlcNAc (**Figure 5B**). This growth defect partially recovered in the later phase of growth, resembling the growth phenotype observed in mucin containing media. These data suggest that *N-*acetyl-D-glucosamine metabolism can bypass PDH function and enable Δ*aceE* growth in mucin media.

### GlcNAc metabolism is required for colonization in neonatal mice

For *E. coli* and other bacteria, *N-*acetyl-D-glucosamine serves as an effective carbon and nitrogen source. *E. coli* actively takes up GlcNAc via the phosphoenolpyruvate-dependent phosphotransferase system (PTS) transporter encoded by *nagE* (31–33). Inside the cell, GlcNAc-6-phosphate is converted to glucosamine-6-phosphate and fructose-6-phosphate by *nagA* and *nagB*, respectively (34). Importantly, NagA deacetylates GlcNAc-6-phosphate to glucosamine-6-phosphate, releasing the acetate group (**Figure 6A**). As supplementation with acetate restores growth of the Δ*aceE* mutant, we hypothesized that acetate released by NagA during GlcNAc metabolism might similarly rescue growth of the Δ*aceE* mutant.To test this, we generated a Δ*nagA* mutant as well as a double mutant lacking both *aceE* and *nagA.* We cultured CRE14, Δ*aceE*, Δ*nagA*, and Δ*aceE* Δ*nagA* in M9 minimal media containing GlcNAc (**Figure 6B**). Again, the Δ*aceE* mutant had slower growth than wild-type in GlcNAc, but reached a similar level of growth as wild type by 24 hours. In contrast, the Δ*nagA* and Δ*aceE* Δ*nagA* mutants were unable to grow at all on GlcNAc as the sole carbon source. Complementation of the Δ*nagA* mutant in GlcNAc was enabled by expressing wild-type *nagBA* under their native promoter, while expressing these alleles in the Δ*aceE* Δ*nagA* led to growth kinetics similar to those of the Δ*aceE* mutant (**Supplemental Figure 5A & B**). Finally, supplementing M9 minimal medium containing glucose with acetate restored growth of the Δ*aceE* and Δ*aceE* Δ*nagA* strains (**Supplemental Figure 5C & D**).

**Figure 6.**
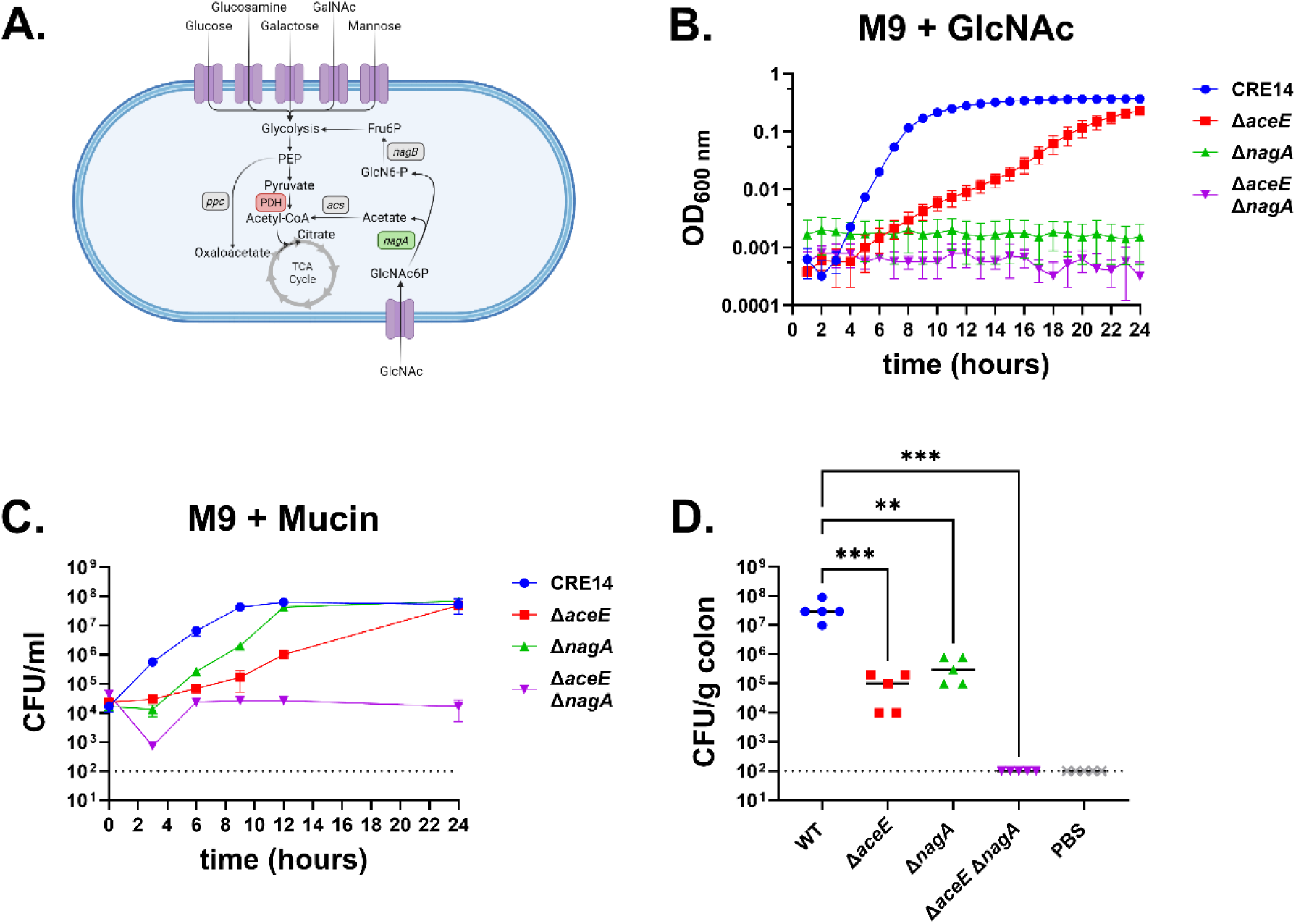
Both the pyruvate dehydrogenase complex and *N-*acetyl-D-glucosamine metabolism are required for growth on mucin and colonization of the infant mouse gut by carbapenem-resistant *E. hormaechei*. A) Schematic of proposed interplay between the TCA cycle and GlcNAc metabolism. Figure created with Biorender.com. B) Growth of CRE14, Δ*aceE*, Δ*nagA*, and Δ*aceE* Δ*nagA* in M9 minimal media supplemented with 10mM *N-*acetyl-D-glucosamine over 24 hours under aerobic conditions. Data represent the mean of three biological replicates and error bars are ± SEM. C) Growth of CRE14, Δ*aceE,* Δ*nagA*, and Δ*aceE* Δ*nagA* in M9 minimal media supplemented with 0.5% partially purified porcine gastric mucin over 24 hours under aerobic conditions. Data represent the mean of three biological replicates and error bars are ± SEM. D) Comparison of colonization abilities of CRE14, Δ*aceE*, Δ*nagA*, and Δ*aceE* Δ*nagA* in the colons of 5-day old infant mice 24 hours post-inoculation with 10^5^ CFU orally (n=5 mice per group). Each point represents the bacterial burden recovered from the colon of an individual mouse, and horizontal bars represent the median CFU/g colon tissue. Statistical significance was determined by One-way ANOVA with multiple comparisons to the WT value. Dashed lines represent the limit of detection.

We then tested if GlcNAc metabolism explained the ΔaceE mutant’s delayed growth on mucin by culturing CRE14, Δ*aceE*, Δ*nagA*, and Δ*aceE* Δ*nagA* in minimal media containing mucin. With mucin as the carbon source, the Δ*nagA* mutant grew better than the Δ*aceE* mutant, but not as well as wild-type. In contrast, the Δ*aceE* Δ*nagA* double mutant was completely unable to grow in mucin media (**Figure 6C**). Taken together, these findings support our hypothesis that acetate released by the action of NagA during GlcNAc metabolism supports growth of the Δ*aceE* mutant on mucin.

To test if mucin metabolism contributes to colonization of the infant gut, we orally administered the four mutant strains to five-day old CD-1 mice and assessed bacterial burden in the colon after 24 hours. The Δ*aceE* and Δ*nagA* mutants colonized to levels 100-fold lower in the colon compared to wild-type, while the Δ*aceE* Δ*nagA* double mutant could not be recovered from the infant mouse colon (**Figure 6D**). These data highlight the significance of both the PDH complex and the GlcNAc utilization pathway on *E. hormaechei* mucin metabolism and colonization of the infant gut.

The function of the bacterial PDH complex is regulated by oxygen (35). Therefore, our findings suggest that an oxygen-dependent metabolic pathway is crucial for population expansion in the gut. The healthy human gut is generally hypoxic due to beta-oxidation by mature intestinal epithelial cells of the short-chain fatty acid butyrate, which is derived from the microbiota (36, 37). This hypoxic environment of the gut lumen limits overgrowth of facultative pathogenic bacteria in the gut. The gut of five-day old infant mice is less hypoxic than that of adult mice due to differences in epithelial cell metabolism (13). We hypothesized that *E. hormaechei* uses oxygen in the gut lumen and metabolizes mucin as a carbon source via the PDH complex.

Growth of carbapenem-resistant *Enterobacteriaceae* such as *E. coli*, *K. pneumoniae*, and *E. hormaechei* in the gut can also be enhanced by broad-spectrum antibiotic treatment, which increases oxygen and availability of nutrients including mucin-derived monosaccharides such as galactose, mannose, and *N-*acetyl-D-glucosamine (23). Our findings show that the oxygen-dependent PDH complex is essential for glucose, glucosamine, galactose, *N-*acetyl-D- galactosamine, and mannose metabolism, whereas the NagA dependent pathway is vital for *N-*acetyl-D-glucosamine metabolism. Both pathways are critical for colonization in the infant gut. This sheds light on mechanisms underlying how carbapenem-resistant *E. hormaechei* may outgrow in the human gut after antibiotic treatment.

### The PDH complex and GlcNAc metabolism facilitate *E. hormaechei* growth in breast milk ***in vivo*.**

Our data suggest that GlcNAc is a major nutrient source for *E. hormaechei* in the gut. Because of this, we questioned if providing additional GlcNAc could enhance *E. hormaechei* gut colonization. The mucin-degrading bacterium *A. muciniphilia* can thrive in human milk and break down human breast milk oligosaccharides (HMOs) due to their structural resemblance to mucin glycans (38). The key components of HMOs include D-glucose, D-galactose, *N*-acetyl*-*D*-* glucosamine (GlcNAc), L-fucose, and *N*-acetylneuraminic acid (39). Despite structural differences between human and mouse milk oligosaccharides, the primary constituents are similar. Thus, we hypothesized that GlcNAc from mouse breast milk could overcome the colonization or growth deficiency of the Δ*aceE* mutant *in vivo*.

To test this, we infected infant mice with wild-type CRE14, Δ*aceE*, Δ*nagA*, and Δ*aceE* Δ*nagA* strains. One group from each infected cohort was returned to their dams, while another group remained separate from the dams. We compared colonization levels between the two groups (housed with or without dams) for each strain (**Figure 7**). Both wild-type and Δ*aceE* colonization increased by approximately 1 log in pups that were returned to dams. We observed colonization defects of similar magnitude in both groups inoculated with the Δ*nagA* strain, indicating that GlcNAc and GlcNAc utilization genes are important for *E. hormaechei* colonization in both cases. Finally, the Δ*aceE* Δ*nagA* double mutant showed similar colonization defects in both groups, suggesting that mouse milk oligosaccharides, particularly GlcNAc, are important carbon sources for *E. hormaechei* colonization.

**Figure 7.**
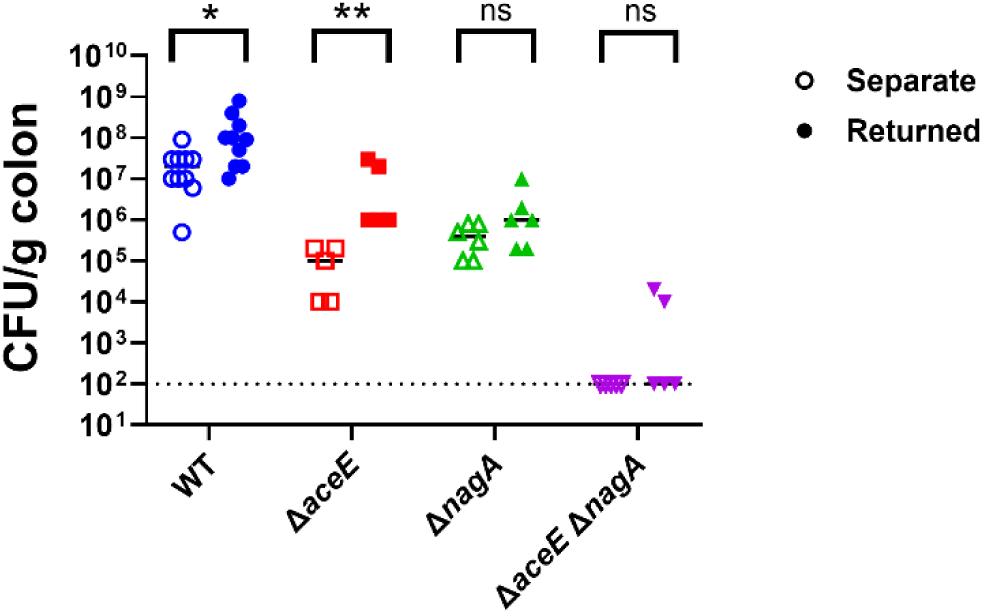
The PDH complex and GlcNAc metabolism aid in utilization of milk oligosaccharides and colonization. Bacterial burden recovered from infant mice inoculated with WT, Δ*aceE*, Δ*nagA*, or Δ*aceE* Δ*nagA* and kept separate from (open symbols) or returned to (closed symbols) dams. Groups of five-day old infant mice were inoculated with 10^5^ CFU of indicated strains and either returned immediately to dams or kept separated. Each point represents the bacterial burden recovered from the colon of an individual mouse 24-hours post-inoculation, and horizontal lines represent the median CFU/g colon tissue from each group. The dashed line indicates the limit of detection. Significance determined by Mann-Whitney nonparametric t-test.

## Conclusions

We provide evidence that *Enterobacter hormaechei* can establish colonization and proliferate in the neonatal mouse gut within 24 hours. *E. hormaechei* primarily colonizes the large intestine of infant mice, likely due to the abundance of mucus present in the colon. These data suggest that *E. hormaechei* can break down mucin and grow on mucin-derived sugars. This is consistent with a previous study that found certain *Proteobacteria*, particularly *Klebsiella*, *Mixta*, *Serratia*, and *Enterobacter* encode several mucin-degrading glycosyl hydrolases (24). We discovered that *E. hormaechei* can use mucin as a carbon source, identifying two critical mucin utilization pathways: the pyruvate dehydrogenase (*aceE)* and N-acetylglucosamine (*nagA*) dependent pathways. Both genes are essential for mucin metabolism and are crucial for colonization and proliferation in the gut.

In our *in vitro* study, we found that *E. hormaechei* CRE14 can utilize GlcNAc, galactose, GalNAc, and mannose as carbon sources. We show the oxygen-regulated pyruvate dehydrogenase complex (*aceE*) is essential for growth in the presence of galactose, GalNAc, and mannose and is required for full colonization of the infant gut. We also found that *nagA* is critical for growth in *N*-acetyl-D-glucosamine, and that *nagA* mutants display a fitness defect in the infant mouse gut, suggesting that *N*-acetyl-D-glucosamine metabolism by *E. hormaechei* is crucial for gut colonization. A mutant lacking both Δ*aceE* Δ*nagA* exhibited complete growth defects in mucin and is incapable of colonizing the infant mouse gut. These findings indicate that these two genes together are crucial for fitness in the gut, making them promising drug targets for developing alternative therapeutic approaches against carbapenem-resistant *E. hormaechei*.

Strategic use of a small animal model, specifically infant mice, to explore the intricate dynamics of *Enterobacter*-host interactions is valuable for studying host-pathogen biology due to its convenience and cost-effectiveness. This model will enable us to efficiently screen mutants, particularly those with defects in colonization during the initial stages of gut colonization. Gaining insights into these early events in *Enterobacter* colonization is important as it offers a foundation for developing effective immunoprophylactic methods to obstruct the initial stages of infection. Understanding and targeting these crucial early interactions will pave the way for more efficient preventative strategies against carbapenem-resistant *Enterobacter* colonization and subsequent infection.

## Methods

### Bacterial strains and culture conditions

*Enterobacter* strains (**Table 1**) were cultured aerobically in LB or M9 minimal medium. Antibiotics were added to agar or broth at the following concentrations: ampicillin (Sigma; 100 µg/ml), apramycin (GoldBio; 50 µg/ml), chloramphenicol (Sigma; 50 µg/ml), and hygromycin B (GoldBio; 100 µg/ml). Partially purified porcine intestinal mucin (Sigma) was added to a final concentration of 0.5%. Human breast milk (Innovative Research, Novi, MI) was added to a final concentration of 50%. Other carbon sources (acetate, glucose, glucosamine, *N-*acetyl-D-glucosamine, galactose, *N-*acetyl-D-galactosamine, fucose, mannose, and sialic acid; Sigma) were added to M9 minimal medium at a concentration of 10 mM.

**Table 1:** Strains.

| Name | Genotype | AtbR | Description | Citation |
| --- | --- | --- | --- | --- |
| ATCC_13047 | WT |  | Wild-type *E. cloacae* ATCC_13047 |  |
| CRE13 | WT | MDR | Wild-type carbapenem-resistant *Enterobacter* spp. UM_CRE-13 | (42) |
| CRE14 | WT | MDR | Wild-type carbapenem-resistant *E. hormaechei* UM_CRE-14 | (42) |
| CRE14/vector | WT<br>pBBR1Hyg | HygR | Wild-type UM_CRE-14 with empty pBBR1Hyg complementation vector | This study |
| $\Delta aceE$ | $\Delta aceE::frt$ | | Isogenic $\Delta aceE$ mutant (PQF87_02045) in UM_CRE-14 | This study |
| $\Delta aceE$/vector | $\Delta aceE::frt$<br>pBBR1Hyg | HygR | Isogenic $\Delta aceE$ mutant with empty pBBR1Hyg complementation vector | This study |
| $\Delta aceE$/paceE | $\Delta aceE::frt$<br>pBBR1Hyg-*pdhR-aceE* | HygR | Isogenic $\Delta aceE$ mutant complemented *in trans* with wild-type *pdhR-aceE* alleles with their native promoter | This study |
| $\Delta nagA$ | $\Delta nagA::frt$ | | Isogenic $\Delta nagA$ mutant (PQF87_04830) in UM_CRE-14 | This study |
| $\Delta nagA$/vector | $\Delta nagA::frt$<br>pBBR1Hyg | HygR | Isogenic $\Delta nagA$ mutant with empty pBBR1Hyg complementation vector | This study |
| $\Delta nagA$/pnagBA | $\Delta nagA::frt$<br>pBBR1Hyg-*nagBA* | HygR | Isogenic $\Delta nagA$ mutant complemented *in trans* with wild-type *nagBA* alleles with their native promoter | This study |
| $\Delta aceE \Delta nagA$ | $\Delta aceE::acc(3)IV$<br>$\Delta nagA::frt$ | ApraR | Double $\Delta aceE \Delta nagA$ mutant in UM_CRE-14 | This study |
| $\Delta aceE$<br>$\Delta nagA$/vector | $\Delta aceE::acc(3)IV$<br>$\Delta nagA::frt$<br>pBBR1Hyg | ApraR<br>HygR | Double $\Delta aceE \Delta nagA$ mutant with empty pBBR1Hyg complementation vector | This study |
| $\Delta aceE$<br>$\Delta nagA$/pnagBA | $\Delta aceE::acc(3)IV$<br>$\Delta nagA::frt$<br>pBBR1Hyg-*nagBA* | ApraR<br>HygR | Double $\Delta aceE \Delta nagA$ mutant complemented *in trans* with wild-type *nagBA* alleles with their native promoter | This study |

### Mutant construction and complementation

A complete list of primers and plasmids can be found in **Supplemental Tables 1 and 2**. Mutant strains were generated by λ Red recombineering. Deletion-insertion alleles were generated by PCR by amplifying the *acc*(*3*)*IV* apramycin resistance cassette from pUC18-miniTn7T-apra with primers containing 50 base pairs of homology to the target gene. The first and last three codons of the open reading frame were preserved. PCR products were introduced into CRE14 harboring pSIM18 by electroporation. Putative recombinants were verified by PCR and all mutant strains were cured of pSIM18. The apramycin resistance gene was subsequently resolved by expressing FLP recombinase from pCP20. pCP20 was transformed into mutant strains by electroporation and propagated at 30°C. A single colony was struck onto plain LB and incubated at 42°C for 6 hours, then transferred to 37°C until the following morning. Resulting colonies were replica patched onto selective media and colonies which were sensitive to apramycin and chloramphenicol were selected. Resolution of the apramycin resistance cassette was also confirmed by PCR. The double Δ*aceE* Δ*nagA* mutant was generated through sequential rounds of λ Red recombineering. Briefly, the Δ*aceE* mutation was generated in a Δ*nagA::frt* background by electroporating the Δ*aceE::acc3(IV)* insertion-deletion fragment into Δ*nagA* harboring pSIM18. Recombinants were confirmed by PCR and the mutant strain was cured of pSIM18 prior to use in phenotypic assays.

The pBBR1Hyg complementation vector was generated by replacing the chloramphenicol acetyltransferase (*cat*) gene on pBBR1MCS with a hygromycin resistance cassette. Briefly, the pBBR1MCS backbone, excluding P*_lac_*, was linearized by PCR. The hygromycin resistance cassette was amplified from pSIM18 with primers containing 20 bp homology to up- and down-stream of the *cat* cassette on pBBR1MCS. pBBR1Hyg was assembled by Gibson Assembly. P*_lac_*was excluded so cloned genes could be expressed from their native promoter. Mutants were complemented with wild-type alleles under expression from their native promoter. *aceE* and *nagA* are both second in an operon, so the upstream gene and promoter region was included in complementation vectors. The pBBR1Hyg plasmid was linearized at the multi-cloning site by PCR with primers. Wild-type *pdhR-aceE* and *nagBA* were amplified from the CRE14 genome with primers containing 20 bp homology to pBBR1Hyg. Complementation vectors were constructed by Gibson Assembly. Resulting colonies were screened by PCR for successful insertion of the desired genes. Complementation vectors were then transformed into indicated mutant strains by electroporation and selection on LB agar containing hygromycin.

### Infant mouse colonization

All animal experiments were approved by the Institutional Animal Care and Use Committee at Michigan State University. Infant mice were infected as previously described. Briefly, five-day-old CD-1 mice (Charles River), orogastrically inoculated with 10^3^ to 10^8^ CFU of indicated bacterial strains. Pups were separated from dams 2 hours prior to inoculation and were either kept at 30°C or returned to dams immediately after inoculation for experiments assessing long-term colonization. Pups were euthanized 24- or 72-hours post-inoculation by isoflurane inhalation. Intestinal segments were aseptically harvested, weighed, and homogenized in 200 µl PBS. Homogenates were serially diluted and plated on LB agar supplemented with ampicillin for CFU enumeration (26, 29).

### Histology

Colon tissues were fixed in 10% neutral-buffered formalin and stored in 60% ethanol. Tissues were sectioned at 5 μm, mounted on frosted glass slides, and stained with hematoxylin and eosin (HE) and PASH stain. Blinded samples were numerically scored for signs of inflammation, such as epithelial damage, inflammatory cell infiltration, goblet cell depletion, cryptic hyperplasia, and cryptic abscess, as follows: 0, none; 1, minimal; 2, mild; 3, moderate; 4, marked; and 5 severe (40).

### CaCo2 adhesion and invasion assays

Caco2 cells were grown in Dulbecco modified Eagle medium (DMEM) supplemented with 10% fetal bovine serum (FBS) and 1% penicillin-streptomycin (Pen-Strep) to confluence, then seeded at 5 × 10^5^ cells per well and allowed to adhere overnight. One hundred microliters of the individual bacterial strains (approx. 1×10^7^ CFU) were added to the appropriate wells and were briefly centrifuged at 3,000 rpm for 1 min. Wells were then incubated in aerobic conditions for 30 min at 37°C. For adhesion assays, the supernatant was aspirated, and the cells were washed three times in 1× PBS and then lysed by incubating in 0.5% Triton X-100 at room temperature for 5 min. Cells were then serially diluted in 1× PBS, plated on LB agar plates, and incubated at 37°C in acerbically for 24 h to enumerate the bacteria. For invasion assays, 100 μl of DMEM containing 10% FBS and 100 μg/ml gentamicin sulfate was added to the cells and allowed to incubate for 1 h at 37°C under aerobic conditions. After incubation, the supernatant was aspirated, and cells were washed three times in 1× PBS and lysed by incubating in 0.5% Triton X-100 at room temperature for 5 min. Cells were then serially diluted and plated on LB agar to enumerate the bacteria inside the cells (19).

### Partial purification of mucin

Porcine gastric mucin from Sigma (Type III) was partially purified as previously described. Briefly, mucin was dissolved in 10% NaCl. The pH was adjusted to 7.0 with NaOH pellets and the mixture was left stirring overnight. Impurities were then separated by centrifugation at 10,000 x g for 10 minutes. The remaining mucin was precipitated by adding ethanol to achieve a final concentration of 60%. After centrifugation to collect the precipitate, the pellet was redissolved in 0.1 M NaCl and precipitated a second time with 60% ethanol. The resulting purified precipitate was then lyophilized and stored at 4°C until further use (40).

### Growth curves (OD)

Overnight cultures of indicated strains were washed twice in sterile PBS and adjusted to an OD_600nm_ of 0.1. Strains were further diluted 1:100 in indicated media for a starting OD_600nm_ of 0.001 in 96 well flat bottom plates and incubated at 37°C under aerobic conditions. Growth was monitored by measuring the optical density at 600nm every 30 minutes for 24 hours under in a BioTek Epoch 2 plate reader. Each biological replicate was processed in technical triplicate. The reported value is the absolute value of the averaged triplicate OD_600nm_ with blanks subtracted to avoid negative values when strains did not grow under a specific condition.

### Growth curves (CFU)

Overnight cultures of indicated strains were washed twice in sterile PBS and adjusted to an OD_600nm_ of 0.01. Strains were further diluted 1:1,000 in M9 containing 0.5% partially purified mucin media for a starting CFU/ml of 10^4^. Strains were incubated at 37°C with shaking for 24 hours under aerobic conditions. At indicated timepoints 100µl aliquots were removed, serially diluted, and plated onto plain LB medium for CFU enumeration.

## Ethical Approval

The Institutional Animal Care and Use Committee at Michigan State University approved all animal experiments (PROTO202100221)

## Acknowledgments

This work was supported by NIH AI134731 and the Rudolph Hugh Endowment at Michigan State University (V.J.D.). We are grateful to Dr. Linda Mansfield for helpful discussions regarding this work, and Dr. Ayush Kumar for providing pUC18-mini-Tn7-apra.

## Contributions

Conceptualization: R.S., E.N.O and V.J.D.

Methodology: R.S., E.N.O.

Investigation: R.S, E.N.O, T.N.

Writing- Original Draft: R.S., E.N.O.

Writing- Review and Editing: R.S., E.N.O, T.N., S.R.S, and V.J.D.

Visualization: E.N.O.

Project Administration: V.J.D. Funding Acquisition: V.J.D.

